# Genetic drift shapes the evolution of a highly dynamic metapopulation

**DOI:** 10.1101/2022.07.10.499462

**Authors:** Pascal Angst, Camille Ameline, Dieter Ebert, Peter D. Fields

## Abstract

The dynamics of extinction and (re)colonization in habitat patches are common features of metapopulations, causing them to evolve differently than large, stable populations. The propagule model, which assumes genetic bottlenecks during colonization, posits that newly founded subpopulations have low genetic diversity and are genetically highly differentiated from each other. Immigration may then increase diversity and decrease differentiation between subpopulations. Thus, older and/or less isolated subpopulations are expected to have higher genetic diversity and less genetic differentiation. We tested this theory using whole-genome pool-sequencing to characterize nucleotide diversity and differentiation in 60 subpopulations of a natural metapopulation of the cyclical parthenogen *Daphnia magna*. For comparison, we characterized diversity in a single, large, stable *D. magna* population. We found reduced (synonymous) genomic diversity, a proxy for effective population size, weak purifying selection, and low rates of adaptive evolution in the metapopulation compared to the large, stable population. These differences suggest that genetic bottlenecks during colonization reduce effective population sizes, which leads to strong genetic drift and reduced selection efficacy in the metapopulation. Consistent with the propagule model, we found lower diversity and increased differentiation in more isolated, younger subpopulations. Our study sheds light on the genomic consequences of extinction–(re)colonization dynamics to an unprecedented degree, giving strong support for the propagule model. We demonstrate that the metapopulation evolves differently from a large, stable population and that the evolutionary process is largely driven by genetic drift.

## Introduction

Metapopulations, i.e., interconnected populations with extinction–(re)colonization dynamics, are ubiquitous (Hanski et al., 2017). They differ from larger, longer-lived populations largely due to this extinction–(re)colonization dynamic, a key evolutionary feature that introduces genetic bottlenecks during the founding of subpopulations (Hanski, 1999; Hanski et al., 2017; Wang & Altermatt, 2019). As suggested by the propagule model (Slatkin, 1977), new subpopulations are founded when empty habitat patches are colonized by one or a few individuals, often originating from a single source population. The genetic bottlenecks can then lead to high genetic differentiation among new subpopulations and low diversity within subpopulations (or low effective population size). These effects, in turn, lead to increased genetic drift and inbreeding (Cosentino et al., 2012; Montero-Pau et al., 2018; Saastamoinen et al., 2018; Tortajada et al., 2009; Waples, 2017). Conversely, gene flow can have the opposite effect in metapopulations (Hanski & Gaggiotti, 2004; Pannell & Charlesworth, 2000): with a continued influx of immigrants, older subpopulations might become genetically more diverse and less differentiated from each other than newly founded subpopulations.

Because common population genetic models were originally developed for large, panmictic populations, extinction–(re)colonization dynamics received little attention in early population genetics. Although the classical island model could be expanded to include migration and its variation among populations (Wright, 1931), it was only much later that Levins (1969) first introduced the concept of a metapopulation, which considered the extinction and (re)colonization of subpopulations (Wang & Altermatt, 2019). While Levins’ model primarily addressed ecological questions, subsequent efforts have sought to discover how metapopulation dynamics affect genetic diversity. For example, recurrent genetic bottlenecks due to high turnover, i.e., frequent extinction and (re)colonization, might result in high genetic differentiation and low genetic diversity (or low effective population size), which needs to be considered in population genetic models (Giles & Goudet, 1997; McCauley, 1989; McCauley et al., 1995; Slatkin, 1977; Wade & McCauley, 1988; Whitlock, 1992; Whitlock & McCauley, 1990).

Ecological factors that contribute to evolution may vary among metapopulations, particularly with regard to the strength of extinction–(re)colonization dynamics (Bonte & Bafort, 2019; Hedrick & Gilpin, 1997; Ingvarsson et al., 1997; Molofsky & Ferdy, 2005; Pannell & Charlesworth, 2000). On one side of the continuum, a metapopulation could experience low extinction–(re)colonization dynamics with large, stable subpopulations, and high gene flow, whereas on the other side, it could exhibit high extinction–(re)colonization dynamics in a setting of small and unstable habitat patches. A metapopulation’s position along this continuum affects the probability of allele fixation (Charlesworth, 2009; Johri et al., 2021; Whitlock, 2003). Specifically, the fixation probability of an emerging allele in a metapopulation depends not only on its selection coefficient, but also on the effective size, *N_e_*, of subpopulations and the degree of population structure (Charlesworth et al., 2003; Vuilleumier et al., 2008; Whitlock, 2003). Variations in population size and structure, as well as their influence on the evolutionary process, can be studied using population genetic summary statistics, such as (non-)synonymous genomic diversity, π_N_ and π_S_, and the rate of (non-)adaptive nonsynonymous substitutions, *ω_NA_* and *ω_A_*. Thus, population genetics can help determine where a particular metapopulation falls along the outlined continuum and give insight into how evolutionary mechanisms differ among metapopulations (Gaggiotti & Foll, 2010).

The position of a given metapopulation along the dynamics-continuum determines the relative importance of natural selection and genetic drift on evolution. Although Hedrick & Gilpin (1997) discussed the factors that influence evolution in metapopulations early on, these issue have not been much addressed in metapopulation genetic studies (but see, e.g., Montero-Pau et al. (2018) for a theoretical model). Metapopulations with stable subpopulations resemble the island model of connected Wright–Fisher populations (Wright, 1931). In these metapopulations, genetic bottlenecks are rare or weak, so evolution is predominantly driven by natural selection (Ronce, 2007; Whitlock, 2004), and local adaptation can help maintain or promote population differentiation and counteract gene flow that would otherwise genetically homogenize subpopulations (Szép et al., 2021). The benthic reef gastropod *Haliotis laevigata* in southern Australia is an example of this kind of metapopulation with large effective population sizes, high connectivity, and low turnover (Sandoval-Castillo et al., 2018).

Strong, frequent bottlenecks can lead to small effective population sizes in which genetic drift predominates (Charlesworth et al., 2003). This process, in turn, weakens natural selection against deleterious mutations and rates of adaptive evolution. The North American Gila Trout (*Oncorhynchus gilae*) shows this pattern of a metapopulation with small effective population sizes, low gene flow, and genomic bottlenecks (Camak et al., 2021). The bottlenecks and the associated low effective population sizes result in inbreeding and accelerate the accumulation and fixation of deleterious mutations, which reduces the mean fitness, referred to as local drift load (Whitlock, 2004). In extreme cases, it results in mutational meltdown of populations (Lynch et al., 1995). Gene flow can counteract this process, introducing new genotypes into a population. In case of hybrid offspring between immigrants and local residents, high drift load can lead to a fitness advantage via hybrid vigor and the subsequent purging of deleterious mutations (Ebert et al., 2002; Whitlock et al., 2000).

Systems with clearly defined subpopulations facilitate the study of metapopulation dynamics and their effect on the evolutionary process. Pond-dwelling organisms occur in distinct water bodies, making population boundaries easy to define. Here, we focus on a pond-dwelling species, the cyclically parthenogenetic microcrustacean *Daphnia magna*, which forms a large metapopulation on the Skerry Islands of southwestern Finland and along the Swedish east coast. As previous findings have suggested, this metapopulation follows the propagule model and is highly dynamic, i.e., characterized by small and unstable subpopulations, high extinction–(re)colonization dynamics, and strong colonization bottlenecks. A long-term survey of this metapopulation (Dubart et al., 2020; Ebert et al., 2013; Pajunen & Pajunen, 2003) has revealed high turnover rates: of the 20 % of the rock pools that contain *D. magna* subpopulations, about 20 % go extinct every year, and about 5 % of the empty ponds are colonized per year. This metapopulation has been shown to be an “inverse mainland-island” type of metapopulation (Altermatt & Ebert, 2010), where the pool of migrants primarily come from small subpopulations. Empty habitat patches are primarily colonized (∼90 % of the time) by single colonizers that then undergo clonal expansion (Haag et al., 2005). While isolated aspects of the system are understood, the question remains whether overall, genetic drift or selection drives evolution in this metapopulation. For example, how does evolution at (non-)synonymous sites in the nuclear genome vary in this metapopulation compared to in large stable populations?

In this *D. magna* metapopulation genomic study, we use allele frequency and ecological data to test our hypotheses about (1) genomic diversity, (2) population differentiation, and (3) (non-)adaptive evolution. Our objective is to understand how this metapopulation evolves and, more generally, how evolution in metapopulations differs from the evolution of larger, longer-lived, stable populations. Considering Haag et al. (2005) who partially explained genetic diversity by a pond’s age and its distance from the sea based on three allozyme markers in the same metapopulation and suggested that the impact of the sea influences the risk of extinction, we revisit the effects of age, ecology, and geography on genomic diversity using whole-genomic data. We expect source sampling under the propagule model in our system to lead to lower genomic diversity in newly established *D. magna* subpopulations than in older ones. Regarding population differentiation, the recurrent genomic bottlenecks might lead to high differentiation between founder populations, which might, however, erode over time due to gene flow (Haag et al., 2005). This high differentiation between founder populations could prevent a pattern of genetic isolation-by-distance (IBD), which is seen in larger scale data for *D. magna* (Fields et al., 2015) across small geographic distances. We further investigate the relative strength of natural selection versus genetic drift using statistics that estimate the proportion of (non-)synonymous polymorphisms like π_N_ and π_S_ and (non-)adaptive substitutions like *ω_NA_* and *ω_A_*. Because of recurring bottlenecks, *N_e_* should be low, which might weaken the efficiency of natural selection and strengthen the effect of genetic drift. Under this scenario, we predict a genomic signature of weak purifying selection, i.e., an excess of nonsynonymous polymorphisms (high ratio of π_N_/π_S_), and few polymorphisms fixed by adaptation (low *ω_A_*). By understanding the variation in genomic diversity, genomic differentiation, and (non-)adaptive evolution in this metapopulation, we try to unravel the general principles of evolution in metapopulations and how they evolve differently from large, stable, panmictic populations using empirical and simulated data.

## Material and Methods

### Study system

This study uses samples from a natural *Daphnia magna* metapopulation located on the Tvaerminne archipelago in southwestern Finland (59°50ʹN, 23°15ʹE). *D. magna* is a Holarctic-distributed, freshwater planktonic crustacean that inhabits small rock pools on the islands of this archipelago. The rock pools (mean volume about 300 L) are depressions in the bare rock of the islands that fill with rainwater but also sometimes collect seawater. We call them rock pools or ponds to avoid confusing them with “pools” from our genomic pool-seq samples. The shallow rock pools are mostly frozen solid in winter; many dry up during the summer as well. Environmental variables including pond geometry, water salinity, humic acid content, pH, calcium concentration, distance to the sea, and height above sea level are available for all ponds (Ebert et al., 2013; Pajunen & Pajunen, 2003; Ranta, 1979). Since 1982, these ponds have been surveyed biannually for the presence of *D. magna*, and since 2007, we have assayed *D. magna* samples from these ponds for parasitic infections (D. Ebert, unpublished data). *Daphnia magna* is a cyclic parthenogen; sexual reproduction results in resting eggs that allow it to survive the winter freezes and the drying out of ponds during the summer. These resting eggs disperse passively by wind, water, or birds and are therefore crucial for migration and colonization of vacant habitat patches. *Daphnia magna* also reproduces asexually, which allows clonal expansion after colonization.

### Samples and sequencing

Our aim was to collect *D. magna* from all occupied ponds (subpopulations) in the core sampling area in late May/early June of 2014 (Figure 1, Table 1 and S1) in order to sequence the pooled genotypes, called pool-seq. Pool-seq is a powerful and cost-efficient way to estimate the genome-wide allele frequencies of populations, as it provides allele frequency estimates of SNPs that are mostly comparable to individual-based sequencing at less cost and with less time (Chen et al., 2022; Dorant et al., 2019; Gautier et al., 2013; Kurland et al., 2019). We collected random subpopulation samples by sieving with hand-held plankton nets through the ponds, aiming for 50 animals per pond and excluding ponds with very small populations at the time of sampling to avoid disrupting natural dynamics. In total, we collected 62 subpopulation samples from 13 islands (Figure 1, Table 1). Collected animals started a three-day regime of antibiotics within 24 hours of collection (Fields et al., 2018) and were fed dextran beads (Sephadex ‘Small’ by Sigma Aldrich: 50 μm diameter) at a concentration of 0.5 g/100 mL to evacuate their gut content and reduce non-target DNA sequencing. Whole animals were then stored in RNAlater (Ambion) and kept at minus 20 °C until we extracted DNA

**Figure 1:**
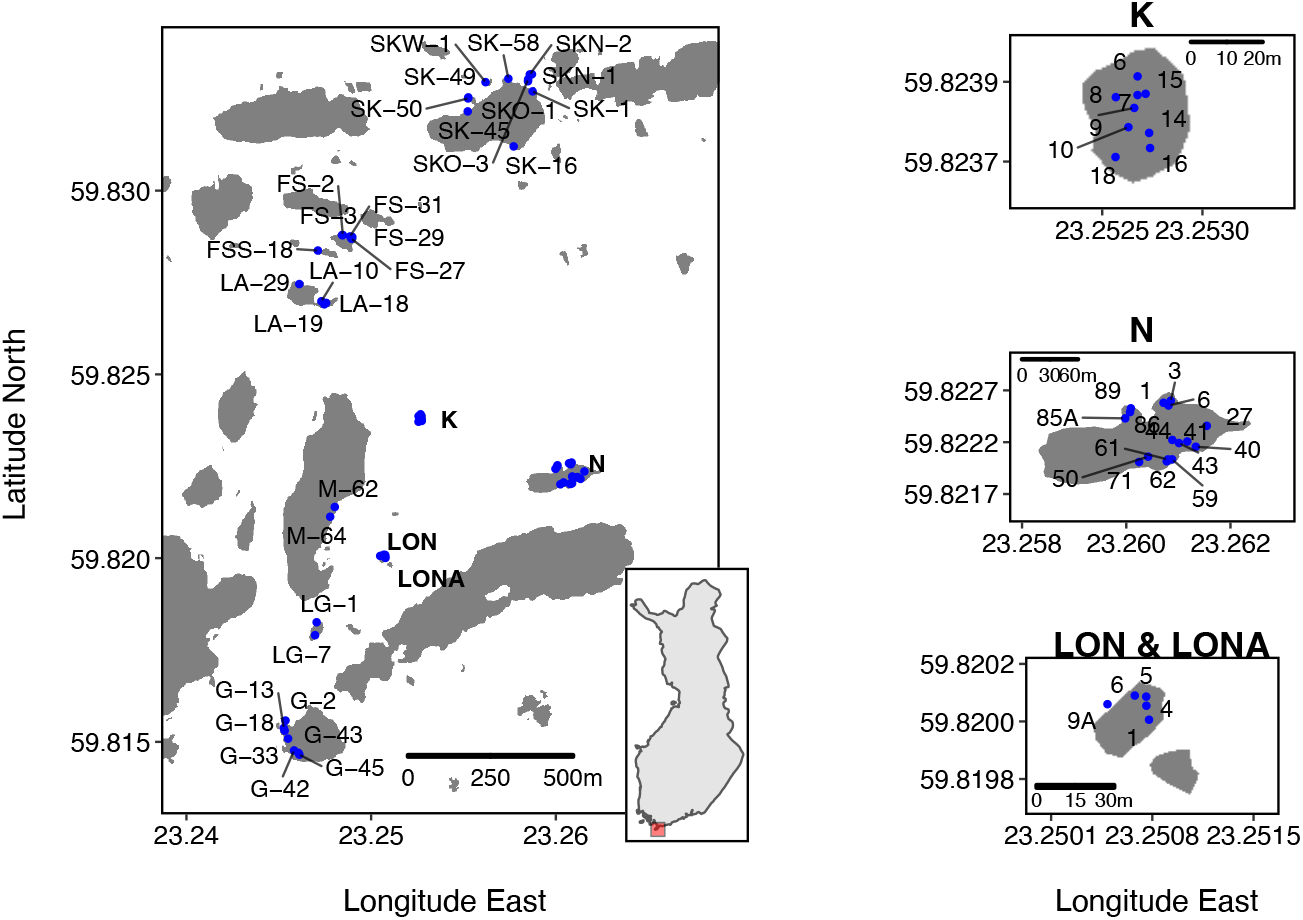
Geographic locations of sampled ponds (subpopulations). Large map shows the overview of the study site, with a small inset map of Finland indicating the location of the study site in red. Islands K, N, and LON/LONA are enlarged in separate maps to improve spatial resolution. Each pond has a unique identifier composed of its island’s ID (see Table 1) and a consecutive number. In the study area are about nine times more habitat patches than shown here. These other habitat patches did not have *D. magna* populations at the time of sampling.

**Table 1:** Island information. Latitude and longitude data are obtained from Google maps. N = Number of sampled ponds (subpopulations); the two samples where the sequencing failed are <u>shown as “-1.”</u>

| Island | Island ID | Latitude/Longitude | N |
| --- | --- | --- | --- |
| Nameless skerry south of Fyrgrundet | FS | 59.8286/23.2485 | 5 |
| Nameless skerry more south of Fyrgrundet | FSS | 59.8283/23.2476 | 1 |
| Granbusken | G | 59.8152/23.2467 | 7(-1) |
| Flatgrund | K | 59.8238/23.2527 | 9 |
| Lasarettet | LA | 59.8273/23.2461 | 4 |
| Nameless skerry | LG | 59.8180/23.2470 | 2 |
| Nameless skerry | LON | 59.8200/23.2506 | 5 |
| Melanskar | M | 59.8215/23.2470 | 2 |
| Storgrundet | N | 59.8221/23.2593 | 16 |
| Skallotholmen | SK | 59.8319/23.2572 | 6(-1) |
| Nameless skerry north of Skallotholmen | SKN | 59.8332/23.2587 | 2 |
| Nameless skerry east of Skallotholmen | SKO | 59.8330/23.2584 | 2 |
| Nameless skerry west of Skallotholmen | SKW | 59.8329/23.2563 | 1 |

For DNA extraction, samples were thawed, the RNAlater was removed, and the samples were washed twice with water. We added 500 μL extraction buffer (Qiagen GenePure DNA Isolation Kit) to the sample tube and ground the sample using a plastic pestle. Then, we added 20 μL Proteinase K for overnight incubation at 55 °C, after which we added 20 μL RNAse for RNA digestion for one hour at 37 °C. For protein removal and DNA precipitation, we followed the instructions on the Qiagen GenePure DNA Isolation Kit, with the addition of 2 μL glycogen (Sigma-Aldrich) to aid DNA precipitation. We then suspended the purified DNA in 80 μL of Qiagen DNA hydration solution and measured DNA concentration using a Qubit 2.0 (Invitrogen). Libraries were prepared using Kapa PCR-free kits and sequenced by the Quantitative Genomics Facility service platform at the Department of Biosystem Science and Engineering (D-BSSE, ETH) Basel, Switzerland, on an Illumina HiSeq 2500 sequencer. Two samples failed this sequencing step, leaving a total of 60 samples for subsequent analyses (G-33 and SK-1; Table 1).

### Ecological covariates and subpopulation age

We summarized ecological covariates (i.e., catchment area, depth, distance to the sea, electrical conductivity, height above the sea, pH, plant cover, submersion time, and surface area) using a principal component analysis (PCA) in R v.4.0.3 (R Core Team, 2020) (see Dubart et al. (2020) for more details). Measures of area and length were log_10_-transformed beforehand. The first two axes of the PCA explained 43.90 % (PC1 25.39 % and PC2 18.51 %) of the variance and were ecologically meaningful. The first principal component described the impact of the sea or “marineness” (e.g., proximity to the sea, water chemistry (salinity, pH level), plant cover (fewer plants closer to the sea)) and represented a gradient from marine to terrestrial ponds. PC2 described geophysical properties independent of the sea (e.g., pond size, depth, catchment area) and represented a gradient from small to large ponds. The age of the subpopulation was assessed using biannual sampling data, with the maximum observed age being 31.5 years, as sampling started in 1982 (Ebert et al., 2013; Pajunen & Pajunen, 2003). A subpopulation was considered newly established if animals were observed after three consecutive visits of seeing no animals, i.e., animals not being seen for more than a year. The chance of a subpopulation remaining undetected for three visits in a row was estimated to be below two percent, as the detection probability of *D. magna* is 0.74 in this survey (Dubart et al., 2020). The subpopulation age was log_10_(age + 1)-transformed for statistical analyses. The geographical distances between ponds were calculated using the R package *geodist* v.0.0.7 (Padgham & Sumner, 2019) and log_10_-transformed for statistical analyses. Infection status with the locally common microsporidian parasite *Hamiltosporidium tvaerminnensis* (Haag et al., 2011), another ecological factor that may explain genomic diversity in the focal metapopulation (Cabalzar et al., 2019), was determined based on field records and on the presence of sequencing reads in our pool-seq samples.

### Mapping genomic reads and variant calling

Raw reads were assessed for quality with FastQC v.0.11.8 (http://www.bioinformatics.babraham.ac.uk/projects/fastqc) and subsequently trimmed to remove low-quality sequence and adapter contamination using the default setting on Trimmomatic v.0.39 (Bolger et al., 2014). The second run of FastQC confirmed successful trimming. These trimmed, paired-end reads were interleaved with seqtk v.1.2 mergepe (https://github.com/lh3/seqtk). The *D. magna* XINB3 individual genome (BioProject ID: PRJNA624896; Fields et al., in prep.) was used as the reference genome when mapping interleaved reads with bwa-mem2 v.2.2.1 (Vasimuddin et al., 2019). Because this reference genome originates from a genotype collected from the same metapopulation, it is closely related to the metapopulation samples in our study. SAMtools v.1.7 (Li et al., 2009) was used to convert SAM files to BAM files, coordinate-sort individual BAM files, and remove unmapped reads. Read groups were added, and duplicates were marked for individual BAM files using the Picard Toolkit v.2.23.9 (Broad Institute, 2019). The average read depth was estimated using SAMtools function depth. INDELs were realigned with GATK v.3.8 RealignerTargetCreator (McKenna et al., 2010; Van der Auwera et al., 2013), which created target intervals, and GATK IndelRealigner. Variants were called using GATK UnifiedGenotyper. These analyses were conducted at the sciCORE (http://scicore.unibas.ch/) scientific computing center at the University of Basel using a snakemake (Mölder et al., 2021) workflow. The VCF file was filtered to include high quality (QUAL > 30, MQ > 40, QD > 2.0, FS < 60) sites with biallelic SNP variants (i.e., excluding INDEL variants) using vcffilter from the C++ library vcflib v.1.0.0_rc2 (Garrison et al., 2021) and VCFtools v.0.1.16 (Danecek et al., 2011). Depth estimates for each sample at each site were recalculated based on the allelic depths using VcfFilterJdk v.1f97a34 (Lindenbaum & Redon, 2018), as GATK includes uninformative reads in the depth estimate but does not include them in the allelic depth estimates. Afterward, we masked genotypes with depth of coverage (DP) less than ten using VCFtools and BCFtools v.1.9 (Danecek et al., 2021), and we masked genotypes with allele depth (AD) of the minor allele equal to 1 using VcfFilterJdk. By applying this minor allele read count filter for each individual sample, we chose a filter that is DP-aware and thus more conservative than the more commonly used minor allele frequency (MAF) filters in pool-seq studies to avoid sequencing errors (Gautier et al., 2013).

### Sequence variation and population genetic analyses

Overall, synonymous, and nonsynonymous genomic diversities were estimated as π, π_S_, and π_N_, respectively, using SNPGenie v.1.0 (Nelson et al., 2015). Because SNPGenie makes calculations per contig, we split the reference FASTA, annotation, and VCF files into individual contigs using *PopGenome* v.2.7.2 (Pfeifer et al., 2014). We converted the split annotation files to GTF format using GffRead v.0.12.1 (Pertea & Pertea, 2020). We used π_S_ as a proxy for effective population size (Leroy et al., 2021) and tested for the association between π_S_ and pond volume (a rough proxy for population size) and the number of mitochondrial haplotypes (a rough proxy for the number of founders and immigrants). We estimated pond volume as a pyramid based on depth and surface area. We estimated the number of mitochondrial haplotypes by reconstructing them from the trimmed sequencing data, starting with mapping interleaved reads to the mitochondrial reference sequence (V3.1; Fields et al., in prep.) using bwa-mem2. We then used the resulting BAM files as input for *RegressHaplo* v.0.1 (Leviyang et al., 2017) in R to reconstruct haplotypes of all samples individually with default parameters. Specifically, we sub-set BAM files using BEDtools v.2.30.0 to focus on a genetic region (from position 10,800 to 12,800) without long conserved regions to avoid performance issues (Leviyang et al., 2017) (Quinlan, 2014). For these calculations made in R, we used the package *data.table* 1.13.6 (Dowle & Srinivasan, 2020) and log_10_-transformed estimates of π_S_, pond volume, and the number of mitochondrial haplotypes. We used the R package *poolfstat* v.2.0.0 (Gautier et al., 2022) to estimate pairwise genomic differentiation based on the VCF file. We removed sites with missing genotypes, converted the pooldata object generated with *poolfstat* to allele frequencies, and then conducted PCA with the *pcadapt* v.4.3.3 (Privé et al., 2020) R package and t-SNE using *Rtsne* v.0.15 (Krijthe, 2015) R package with five dimensions retained from the initial PCA step, perplexity 19, and 5,000 iterations. For generating the input for GESTE v.2 (Foll & Gaggiotti, 2006), which is a Bayesian method based on the *F*-model to estimate *F*_ST_ of subpopulations and to relate *F*_ST_ to environmental factors using a generalized linear model, we used *poolfstat*’s function pooldata2genobaypass() to convert the VCF file to allele read counts. After conversion, we corrected for the corresponding haploid pool sizes as described in Feder et al. (2012) using a modified version of the script baypass2bayescan.py (Stern & Lee, 2020).

### Associations between covariates, genomic diversity, and genomic differentiation

To find associations between genomic diversity and ecological covariates (i.e., subpopulation age, PC1, PC2, mean distance to the two nearest neighbors (NN2), *H. tvaerminnensis* infection status), we performed a multiple regression analysis using a type two ANOVA from the *car* v.3.0-10 (Fox & Sanford, 2019) R package. Additionally, after observing island-specific clusters in the dimensionality-reduction analysis of our genomic data, we also included island of origin as a factor in a second model. We ran the Bayesian GESTE methodology with the same covariates and separately checked for associations between pairwise genomic differentiation and each ecological covariate (i.e., geographic distance, mean subpopulation age, mean PC1, mean PC2, and mean NN2) using distance-based Moran’s eigenvector maps (dbMEM) analysis by redundancy analysis (RDA) to test for isolation-by-distance (IBD), isolation-by-environment, or age-specific genomic differentiation. RDAs were performed on the overall data and separately on data from each island. Specifically, we separately transformed each explanatory variable into dbMEMs using the R package *adespatial* v.0.3-14 (Dray et al., 2021) and decomposed the response variable, pairwise *F*_ST_, into principal components using the R base *stats* function prcomp(). The RDAs were done in R using the package *vegan* v.2.5-7 (Oksanen et al., 2020), with significance assessed by 1,000 permutations.

### D. sinensis genome annotation

To estimate (non-)synonymous divergence, d_N_ and d_S_, and the rate of (non-)adaptive nonsynonymous substitutions, *ω_NA_,* and *ω_A_*, we downloaded the available *D. sinensis* genome (ASM1316709v1; GenBank accession: GCA_013167095.1) from NCBI and RNA-seq reads (run accessions: SRR10389290, SRR10389293, and SRR10389294) from EMBL. *Daphnia sinensis* is closely related to *D. magna* (Cornetti et al., 2019). We removed adapters from the reads using fastp v.0.20.0 (Chen et al., 2018) and checked its success with FastQC. To annotate the genome, we used MAKER2 v.2.31.10 (Holt & Yandell, 2011). Specifically, we created a database from the *D. sinensis* genome and individually aligned the trimmed reads to it using STAR v.2.7.4a (Dobin et al., 2013). The database was used to generate reference-assisted transcriptomes with Trinity v.2.12.0 (Grabherr et al., 2011). We checked biological completeness using BUSCO v.3.0.2 (Seppey et al., 2019) and the arthropoda_odb9 gene set (Creation date: 2017-02-07). Whereas we obtained target proteins by applying TransDecoder v.5.5.0 (https://github.com/TransDecoder/TransDecoder) on the transcriptomes, we used diamond v.2.0.11.149 (Buchfink et al., 2021) as well as hmmer v.3.3.2 (hmmer.org) to find matches of the ORFs to swissprot and pfam. Pyfasta v.0.5.2 (https://github.com/brentp/pyfasta/) was used to split intermediate FASTA file outputs. To obtain transcript hints, the individual transcript files were concatenated and mapped using minimap2 v.2.22-r1105 (Li, 2018) before collapsing isoforms using collapse_isoforms_by_sam.py from the Cupcake tool (https://github.com/Magdoll/cDNA_Cupcake).

### Summary statistics of the divergence data

We repeated the methodology for read mapping and VCF file preparation using the *D. sinensis* genome as a reference and estimated (non-)synonymous divergence based on each sample’s average distance to the reference with SNPGenie. The rate of adaptive substitution, *α*, was calculated based on the per-site counts of (non-)synonymous polymorphisms and substitutions (Charlesworth, 1994). Afterward, we calculated *ω_A_* as *α*(d_N_/d_S_) and *ω_NA_* as (1-*α*)(d_N_/d_S_). We tested whether *ω_A_* and *ω_NA_* were correlated with *N_e_* using Spearman correlations. *N_e_* was approximated with genomic diversity at synonymous sites, π_S_, using SNPGenie.

### Empirical data of single, large, stable population

To compare our focal metapopulation with a single, larger, more stable *D. magna* population, we estimated overall genomic diversity, π, genomic diversity at (non-)synonymous positions, π_N_, and π_S_, (non-)synonymous divergence, d_N_ and d_S_, and the rate of (non-)adaptive nonsynonymous substitutions, *ω_NA_* and *ω_A_*, for the *D. magna* population from pond Aegelsee near Frauenfeld, Switzerland (47°33’28.0”N, 8°51’46.0”E; surface area around 30,000 m^2^). This population is at least 60 years old and has an estimated minimum population size of over ten million individuals. The Aegelsee does not entirely freeze in winter nor dry up in summer. However, fall and winter conditions result in little to no overwintering of *D. magna* (Ameline, Bourgeois, et al., 2021). In spring, *D. magna* hatches from resting eggs (Ameline, Vögtli, et al., 2021). We collected a sample of 102 individuals in the spring of 2017, pool-sequenced them and prepared them for analysis identically to the metapopulation samples. To calculate the genomic summary statistics, we used SNPGenie with two separate reference genomes, i.e., *D. magna* and *D. sinensis*.

### Simulations

To further investigate relationships between (non-)synonymous genomic diversity and effective population size in this metapopulation, we simulated different-sized populations and compared their variation in (non-)synonymous genomic diversity with estimates from our collected natural subpopulations. Using a non-Wright–Fisher model in SLiM v.3.6.0 (Haller & Messer, 2019), we simulated a 100 kilobase pair stretch of DNA in different-sized panmictic populations with a recombination rate of 1 × 10^-8^ and a mutation rate of 1 × 10^-8^. We ran the simulations until populations reached mutation-drift equilibrium, which is approximately ten times as many generations as the size of the population (Haller & Messer, 2019). We made simulations with different distributions of the selection coefficient, i.e., distribution of fitness effects (DFE), for nonsynonymous mutations for each population size. The DFE was either fixed at zero or drawn from a gamma distribution. However, we fixed the selection coefficient of synonymous mutations at zero for all runs. We conducted 1,000 replicate simulations for each setting.

To compare the observed (non-)synonymous substitution rates in the metapopulation to in-silico data, we performed a second set of simulations with substitution tracking enabled. We increased the number of generations per simulation to 100,000 and the simulated sequence length to one Mbp to get a significant number of polymorphisms that would reach fixation. Moreover, 15 % of the nonsynonymous mutations were beneficial (s = 0.0001), making the calculation of *ω_A_* and *ω_NA_* more meaningful.

## Results

### Sequencing and population structure

A total of 60 *D. magna* subpopulations distributed throughout almost the entire survey area of the metapopulation were successfully sequenced (Figure 1, Table 1). Between 72 and 99 % of our pool-seq reads were mapped to the *D. magna* reference genome (Table S1). Samples with a low mapping percentage were infected with a locally common microsporidian parasite (*Hamiltosporidium tvaerminnensis*; Table S1) that can dominate the sequencing reads when whole genomes of host and parasites are co-sequenced (Angst et al., 2022). With only a few exceptions, the average read coverage for most *D. magna* samples was beyond 20× (Table S1). After variant filtration, we were left with 1,540,716 SNPs to analyze. A PCA of this SNP data indicated an overall structure in the data (Figure 2A). As expected, most samples originating from the same island clustered together. The PCs reflected geographic distribution to some degree, with longitude being positively correlated with PC1 (Spearman’s *r*(58) = 0.71, *p* < 0.001) and latitude negatively correlated with PC3 (Spearman’s *r*(58) = −0.50, *p* < 0.001). The clustering by island became more defined when samples were pinpointed on a two-dimensional map based on the first five PCs using t-distributed stochastic neighbor embedding (t-SNE; Figure 2B). Interestingly, in some cases, samples from some islands fell into multiple subclusters (LA, M, N) reflecting distinct regions on the islands (Figure S1). These subclusters may represent different colonization histories for these regions. Furthermore, samples from island SK and its close neighboring islands, SKN, SKO, and SKW, formed one cluster, so did samples from three geographically close islands FS, FSS, and LA (Figure 2B).

**Figure 2:**
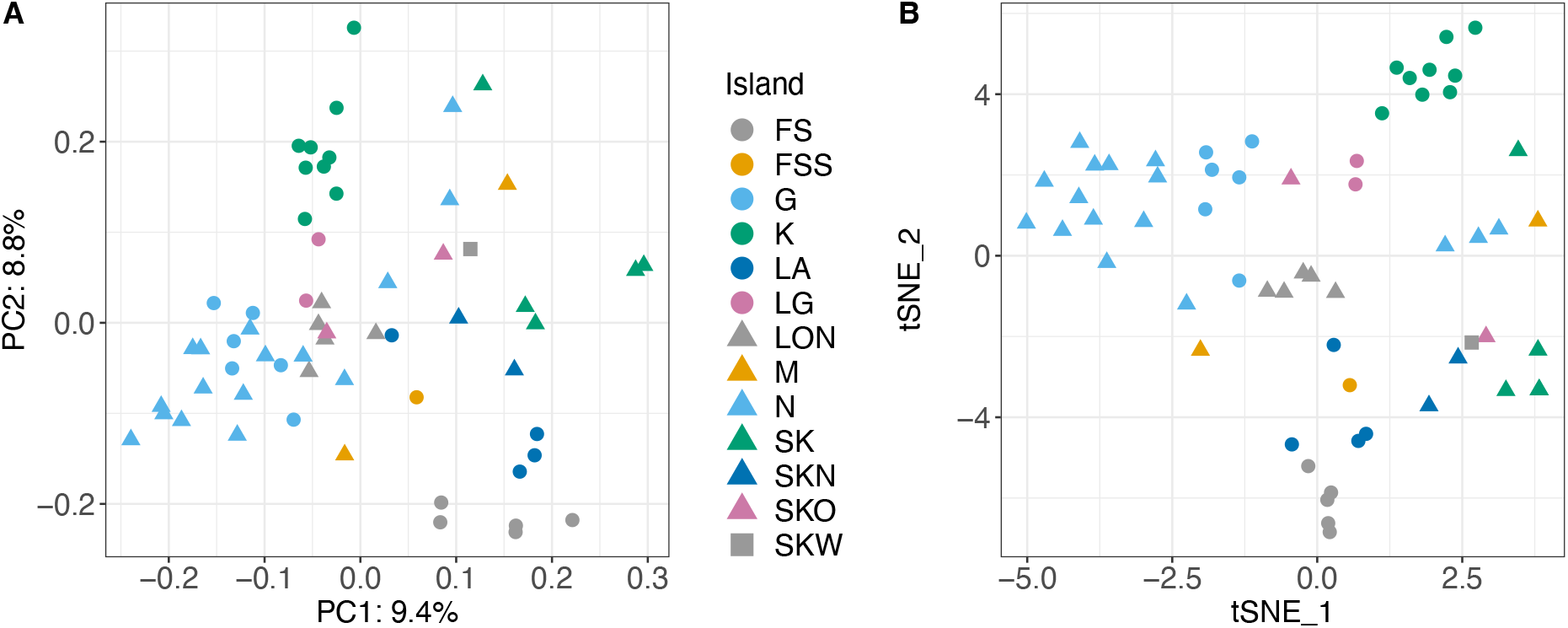
Dimensionality-reduction using PCA and t-SNE. PCA (A) and t-SNE (B) reveal spatial population structure based on whole-genome allele frequency data. Samples from the same island usually form clusters. Colored symbols represent island of origin (see legend). Percentages in A give the amount of variance explained by PC1 and PC2.

### (Non-)adaptive genomic divergence

Synonymous genomic diversity, π_S_, is often used to approximate the theoretical quantity *N_e_*, acting as a stand-in to predict how populations will behave evolutionarily. We tested this approximation by correlating π_S_ with pond size (assuming larger ponds have larger populations and receive immigrants more likely) and with the number of mitochondrial haplotypes (assuming the number of haplotypes is representative of the number of colonists and immigrants). While both variables showed a positive association, as expected, it was not a strong correlation: the correlation of π_S_ with pond size was Pearson’s *r*(58) = 0.25, *p* = 0.05, whereas the correlation of π_S_ with mitochondrial haplotypes was *F*(1,58) = 4.502, *p* = 0.038, and *R*^2^ = 0.07. This pattern is not entirely unexpected because the degree to which *N_e_* correlates with population census size, *N*, is highly variable and depends on a range of ecological and evolutionary details in an individual biological system (Waples, 2022). For example, in this metapopulation, large subpopulations can be founded by one or several individuals that have undergone clonal expansion. Observed nonsynonymous and synonymous genomic diversity, π_N_ and π_S_, were strongly positively correlated but with a slope of less than one (Figures 3A and S2). More diverse subpopulations showed larger departures from a one-to-one ratio, meaning that π_N_ decreased relative to π_S_ as diversity of subpopulations increased (Figures 3A and S2). We found the same relationship between nonsynonymous and synonymous diversity when we drew nonsynonymous mutations from a distribution of negative selection coefficients to simulate populations of different sizes (Figure 3B). When the selection coefficient of all mutations is zero, the slope is one (Figure 3B). Therefore, observed and simulated results coincided with expectations from population genetic theory, that purifying selection is more efficient in removing deleterious nonsynonymous polymorphisms from populations with higher synonymous diversity (or larger populations) (Charlesworth, 2009). Furthermore, the ratio of nonsynonymous to synonymous genomic diversity in the subpopulations, π_N_/π_S_, was positively correlated with the isolation measure (mean distance to the two closest neighboring subpopulations, or NN2) (Spearman’s *r*(58) = 0.36, *p* = 0.005; Figure 5), suggesting that purifying selection is less efficient in more isolated (= less diverse) subpopulations. Positive selection, measured as the rate of adaptive nonsynonymous substitutions, *ω_A_*, appeared higher in populations with higher synonymous diversity, a pattern also seen in the data from the metapopulation estimated with the outgroup *D. sinensis* (Figure 4A, Supplemental Results) and in the simulation results (Figure 4B). Because natural selection and genetic drift are non-independent, positive selection may be more efficient, as it is less affected by genetic drift. (The rate of nonadaptive nonsynonymous substitutions, *ω_NA_*, is an inversion of *ω_A_* and was lower in larger populations (Figures S3 and S4)).

**Figure 3:**
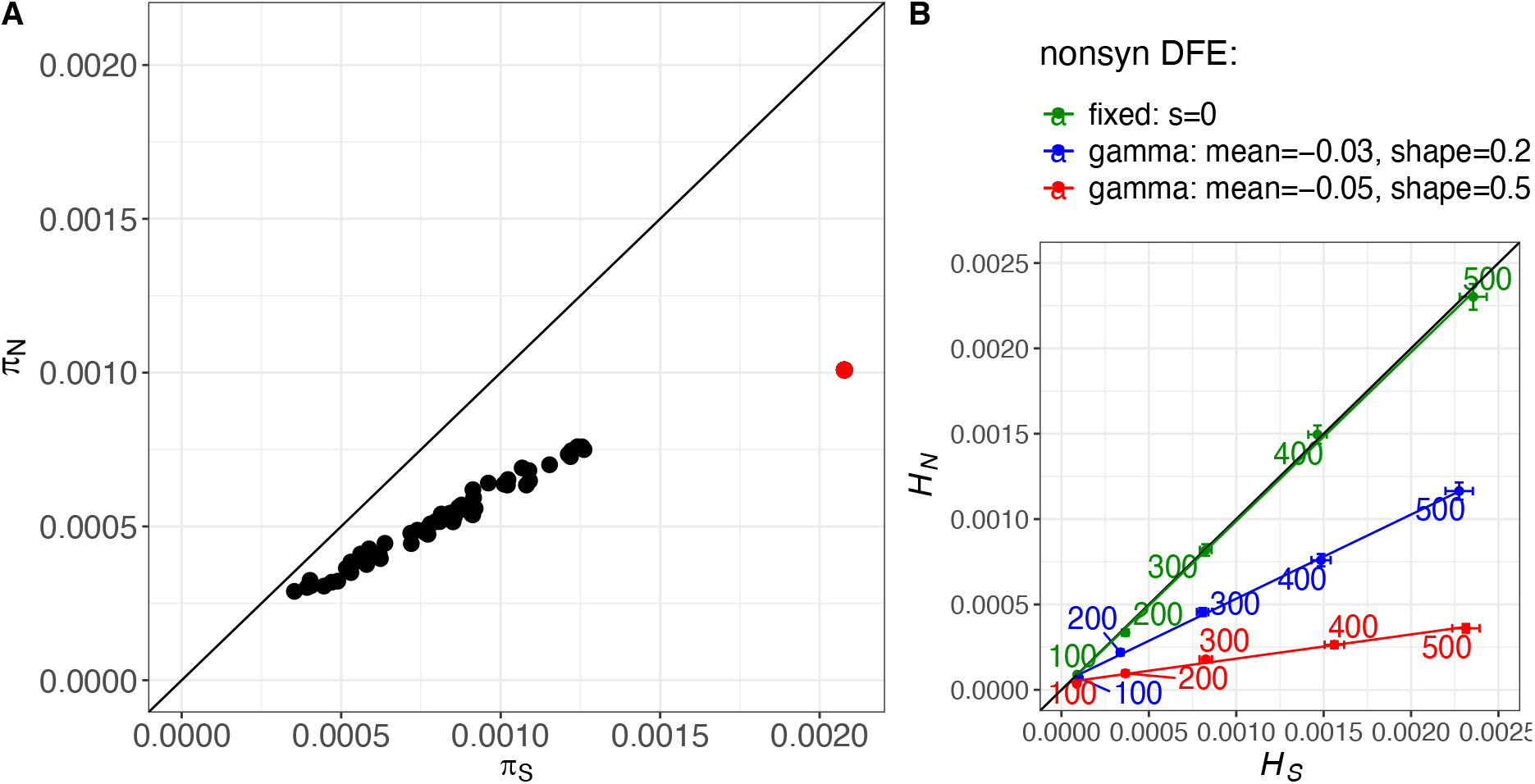
Association between nonsynonymous and synonymous genomic diversity in populations of different sizes. (A) π_N_ plotted against π_S_, with each black dot representing one subpopulation. The slope of the data cloud is smaller than one (0.52). Thus, more diverse subpopulations have relatively fewer nonsynonymous polymorphisms, suggesting that purifying selection is stronger in these subpopulations. The red dot is the estimate for the single, large, stable population from Switzerland. (B) Simulated (non-)synonymous heterozygosities using SLiM (Haller & Messer, 2019). The colored numbers show the simulated population sizes. *H_S_* correlates well with *N_e_*. Colors indicate the nonsynonymous distribution of fitness effects (DFE) following different gamma distributions (blue and red) or fixed to zero (green, total absence of selection; s = 0). Red shows the strongest selection. Error bars indicate the standard error around the mean of 1,000 runs. Colored lines are fits for the different nonsynonymous DFEs using linear regressions. The black line indicates the one-to-one ratio in both plots.

**Figure 4:**
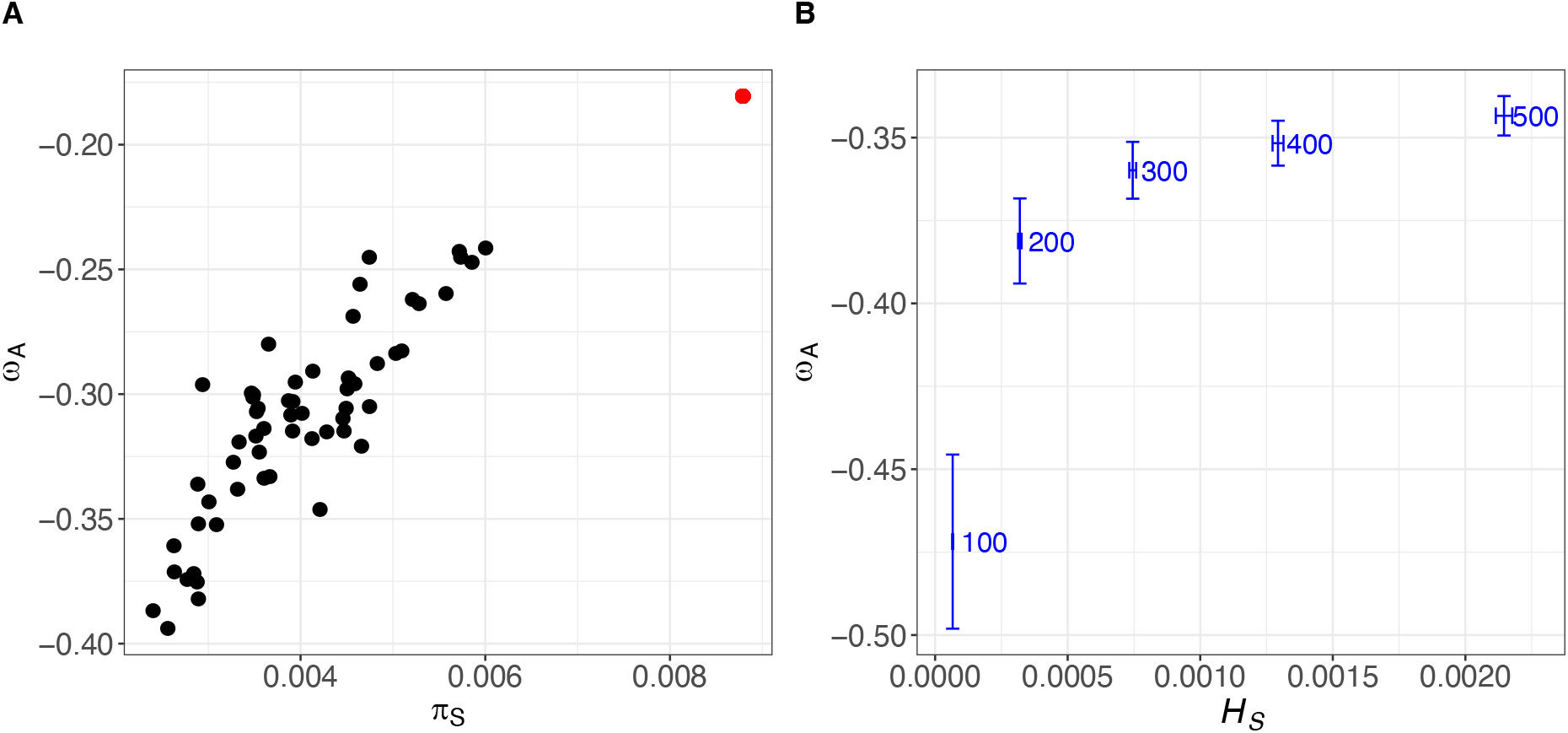
Association between the proportion of adaptive nonsynonymous substitutions, *ω_A_*, and synonymous genomic diversity, π_S_ and *H_S_*, in differently sized populations. (A) Observed *ω_A_* for each subpopulation; *ω_A_* is positively associated with π_S_, which is a useful approximation for *N_e_*. The red dot shows the estimate for the single, large, stable population from Switzerland. (B) Shows the positive association of *ω_A_* with *H_S_* and the simulated population size. Horizontal and vertical error bars indicate the standard error around the mean of 1,000 runs (simulations were performed with SLiM (Haller & Messer, 2019)).

**Figure 5:**
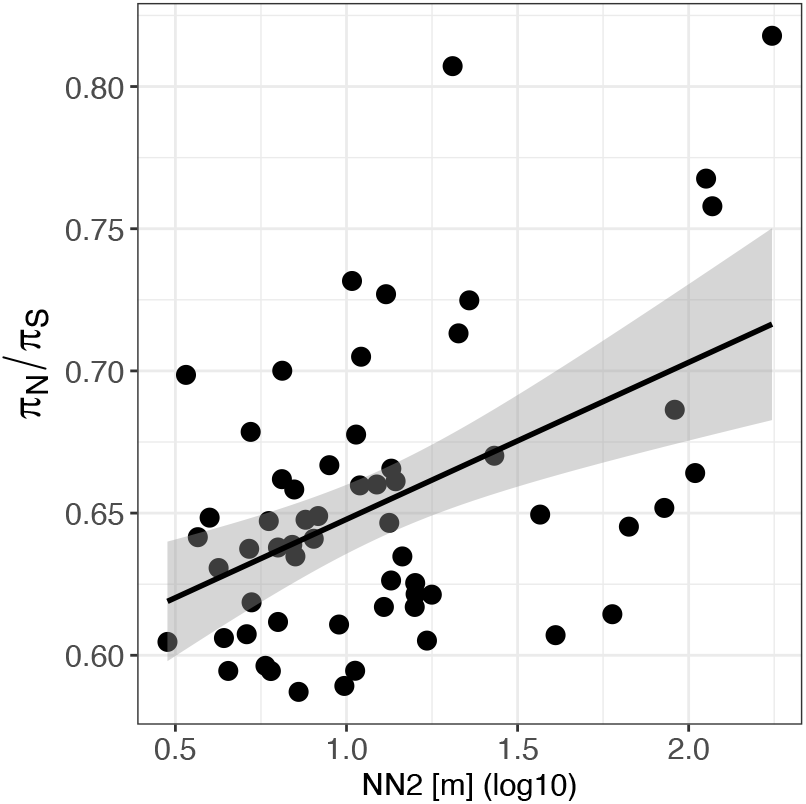
π_N_/π_S_ depends on the degree of spatial isolation of a subpopulation. The number of nonsynonymous polymorphisms relative to synonymous polymorphisms correlates positively with the isolation measure, NN2 [m]. Statistics derive from a Spearman correlation test. Each dot represents one subpopulation. The black line is a regression line from a linear model with the 95 % confidence interval depicted as shading around the line (*R* = 0.36, *p* = 0.005).

### Genomic diversity in relation to ecology

Genomic diversity, π, was estimated separately for each *D. magna* subpopulation. It ranged by nearly an order of magnitude from 3 × 10^-4^ to 1 × 10^-3^ with a mean of 6 × 10^-4^. We tested whether genomic diversity was associated with subpopulation age since colonization, NN2, ecological variables, and infection status. PCA summarized ecological variables, with PC1 representing a gradient from marine to terrestrial ponds, and PC2 representing a gradient from small to large ponds. Using multiple regression analysis, we found that young, isolated subpopulations were less diverse than older, less isolated subpopulations (Table 2, Figures 6 and S5). These findings held true after correcting for the island of origin (Table S2). We found no significant association between π and either PC1 (marineness), PC2 (pond geometry), or *H. tvaerminnensis* infection status in the overall model (Table 2).

**Figure 6:**
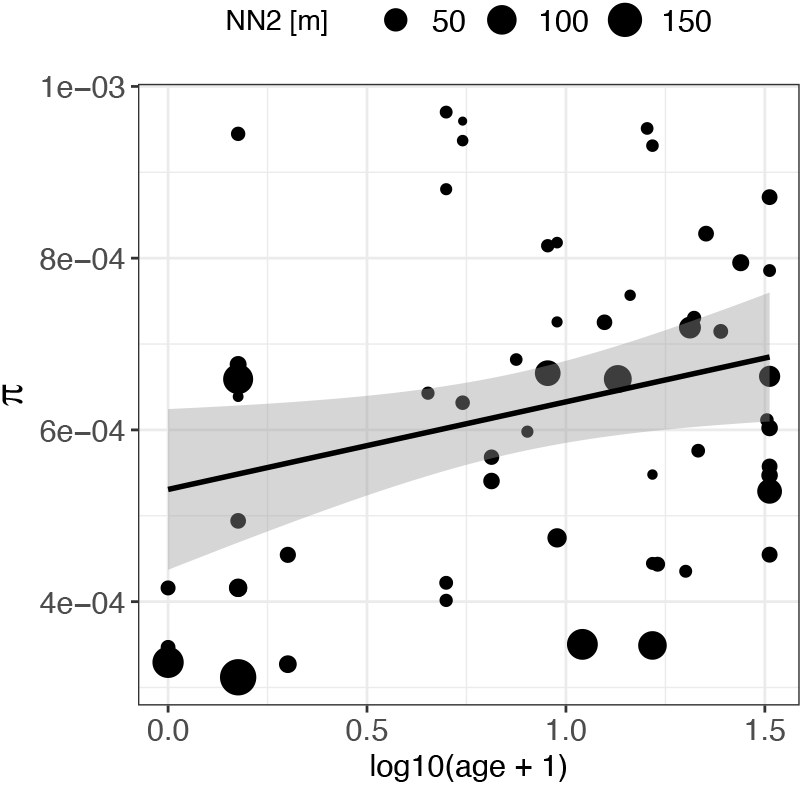
Genomic diversity, π, explained by subpopulation age (x-axis) and isolation level (NN2, size of symbol). Genomic diversity within subpopulations is positively correlated with subpopulation age (positive slope of regression line) and negatively correlated with the isolation measure, NN2 [m] (symbol sizes become smaller towards the top of the graph) (compare Table 2). Population age is log_10_(age + 1)-transformed. Each dot represents a subpopulation. The regression line is based on a linear model (π log_10_(age + 1)) with the confidence interval depicted as shading around the line.

**Table 2:**
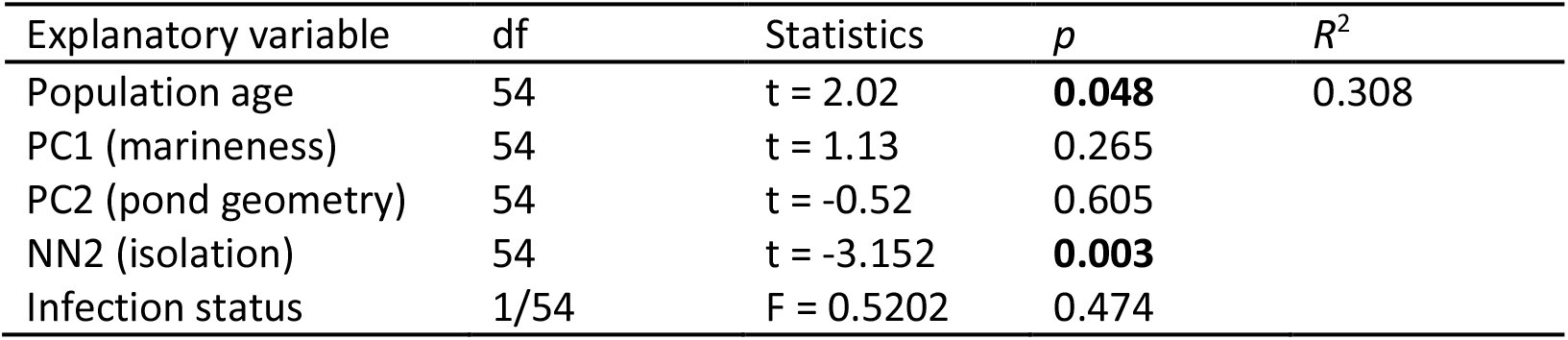
Type two analysis of variance between genomic diversity, π, and explanatory variables. This type of analysis of variance follows the principle of marginality, testing each term after all others (partial sum of squares model). One R^2^ value is given for the complete model. Significant associations are in bold.

| Explanatory variable | df | Statistics | $p$ | $R^2$ |
| --- | --- | --- | --- | --- |
| Population age | 54 | t = 2.02 | **0.048** | 0.308 |
| PC1 (marineness) | 54 | t = 1.13 | 0.265 |  |
| PC2 (pond geometry) | 54 | t = -0.52 | 0.605 |  |
| NN2 (isolation) | 54 | t = -3.152 | **0.003** |  |
| Infection status | 1/54 | F = 0.5202 | 0.474 |  |

### Genomic differentiation in relation to ecology

Genomic differentiation between subpopulations, as estimated with *F*-model-based *F*_ST_ using GESTE, had a mean of 0.06 and a range from 0.02 to 0.21. GESTE’s test for association with subpopulation age, environment, and isolation suggested that genomic differentiation was best explained by a model with only the measure of isolation, NN2, i.e., the model with the highest posterior probability (postProb = 0.48; Table 3, Figures 7A and S6). Models without this isolation measure had a posterior probability of essentially zero, rendering them unlikely. The relationship between *F*_ST_ and NN2 remained significant when we corrected for island of origin and infection status (*F*(1,45) = 67.961, *p* < 0.001, and *R*^2^ = 0.75). Furthermore, as estimated using *poolfstat*, pairwise genomic differentiation correlated with mean NN2 (dbMEM analysis by RDA: *R*^2^ = 0.05, *p* = 0.013; Figure S7). Therefore, both the *F*-model and the pairwise approach for estimating genomic differentiation found that geographically isolated subpopulations showed greater differentiation than less isolated subpopulations, and that geographical isolation of subpopulations was the primary driver among the factors tested for overall patterns of genomic differentiation in the focal metapopulation. Additionally, pairwise genomic differentiation and geographic distance were positively correlated (*R*^2^ = 0.39, *p* = 0.001; Figure 7B), meaning that subpopulations separated by greater geographical distance were more differentiated than geographically closer subpopulations, i.e., isolation-by-distance (IBD). We also found IBD when looking specifically at combinations of subpopulations within islands (mean *R*^2^ = 0.38). Correlations between pairwise genomic differentiation and the remaining candidate variables (PC1 (marineness), PC2 (pond geometry), and subpopulation age) were insignificant (Table 2). This pattern corresponds with results from the GESTE, pointing to IBD as the dominant pattern. On the other hand, the immediate effect of genetic bottlenecks during population founding on higher population differentiation among young subpopulations is not as strong in the overall analysis. However, the finding by Haag et al. (2005) that newly founded subpopulations are more differentiated from each other than older subpopulations was confirmed when looking specifically at comparisons between the age class of newly founded subpopulations and older age classes (<= 2 years old: mean *F*_ST_ = 0.49, CI95 = 0.47-0.51; > 2 and <= 15 years old: mean *F*_ST_ = 0.35, CI95 = 0.33-0.37; > 15 years old: mean *F*_ST_ = 0.36, CI95 = 0.35-0.38; Wilcoxon’s W = 21,774, p < 0.001 and W = 22,103, p < 0.001 for old and intermediate age classes versus young age class, respectively). The finding that intermediate and old age classes did not differ in mean pairwise *F*_ST_ (Wilcoxon’s W = 28,697, *p* = 0.16) reflects the nonsignificant correlation between pairwise population differentiation and mean age (see RDA).

**Figure 7:**
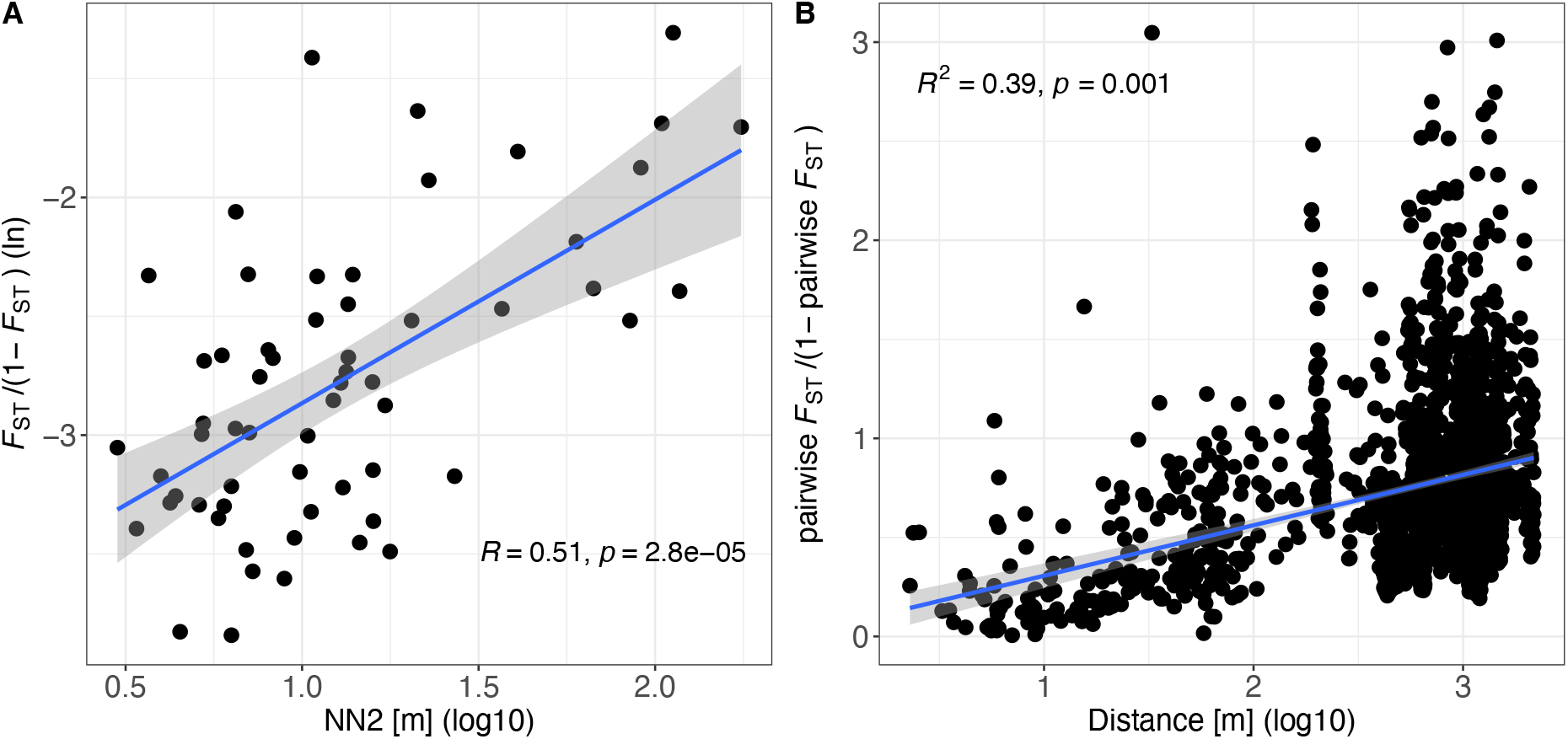
Isolation by distance (IBD). (A) *F*-model-based *F*_ST_ positively correlates with the isolation measure NN2 [m]. Each point represents one subpopulation. Statistics derive from a Spearman correlation test. (B) Pairwise population differentiation correlates positively with geographic distance. Each point represents the comparison between two subpopulations. Statistics derive from dbMEM analysis by RDA. In both plots, the blue line is a regression line based on a linear model with the confidence interval depicted as shading around the line.

**Table 3:** Associations between differentiation in subpopulations, *F*ST, and explanatory variables calculated using GESTE. GESTE is a Bayesian method based on the *F*-model for estimating the *F*ST of subpopulations and for testing associations between *F*ST and explanatory variables. Here, age, PC1, PC2, and NN2 are explanatory variables. Each combination of explanatory variables in the first column is accompanied by a posterior probability in the second column. Unrounded posterior probabilities sum to one. Models without NN2 result in a posterior probability of 0.

| Explanatory variable(s) | postProb(model) |
| --- | --- |
| Constant | 0.000 |
| Constant, age | 0.000 |
| Constant, PC1 | 0.000 |
| Constant, PC1, age | 0.000 |
| Constant, PC2 | 0.000 |
| Constant, PC2, age | 0.000 |
| Constant, PC2, PC1 | 0.000 |
| Constant, PC2, PC1, age | 0.000 |
| Constant, NN2 | 0.478 |
| Constant, NN2, age | 0.012 |
| Constant, NN2, PC1 | 0.027 |
| Constant, NN2, PC1, age | 0.002 |
| Constant, NN2, PC2 | 0.374 |
| Constant, NN2, PC2, age | 0.001 |
| Constant, NN2, PC2, PC1 | 0.094 |
| Constant, NN2, PC2, PC1, age | 0.003 |

### Single, large, stable population

To compare our focal metapopulation’s genetic summary statistics with a single large population of the same species, we used pool-seq data from the relatively large Aegelsee *D. magna* population in Switzerland generated in the same way as for our metapopulation. Based on the analysis of 1,056,626 SNPs, we estimated a π value of 1.5 × 10^-3^, about 2.5 times higher than estimated for the entire metapopulation. The ratio of π_N_ (1.0 × 10^-3^) and π_S_ (2.0 × 10^-3^) corresponded with our expectations for a larger population, as it showed higher π_S_ and more deviation from the one-to-one line than the metapopulation samples (Figure 3A). Also, *ω_A_* (−0.18) and *ω_NA_* (0.32) were higher and lower than in the metapopulation samples, respectively, as expected for a larger population (Figure 4 and S3).

## Discussion

Extinction–(re)colonization dynamics are key in distinguishing metapopulations from larger, more stable populations with gene flow (Hanski, 1999; Wang & Altermatt, 2019). Understanding how these metapopulation dynamics influence molecular evolution is a step toward understanding how metapopulations differ from the more extensively studied Wright–Fisher populations. This study presents evidence that nondeterministic processes play a large role in a highly dynamic, natural metapopulation, and that subpopulation founding with initial genetic bottlenecks leads to subsequent inbreeding and low genomic diversity as assumed under the propagule model. This model predicts strong genetic drift, a weakened efficacy of selection, and the accumulation of deleterious mutations, all of which we observed in the here studied metapopulation. Contrasting this metapopulation with a stable, large population of the same species shows that genomic diversity is considerably lower and genetic drift much stronger in the metapopulation.

### Evolutionary model

By identifying the genomic variation, age, ecology, and geography of individual subpopulations, we investigated the evolution of interconnected populations. Previous research in our focal *D. magna* metapopulation has found high turnover dynamics, small numbers of subpopulation founders, and the accumulation of deleterious mutations in the mitochondrial genome, all consistent with the propagule model (Altermatt & Ebert, 2010; Dubart et al., 2020; Ebert et al., 2013; Fields et al., 2018; Zumbrunn, 2011). Subpopulations tend to be short-lived, undergoing frequent, strong genetic bottlenecks during the colonization of empty habitat patches and subsequently suffering from inbreeding. The effective population size of subpopulations is correspondingly low (Walser & Haag, 2012), and genetic drift is a strong evolutionary force that can reduce natural selection efficacy. Our study corroborated these features, demonstrating low genomic diversity and high differentiation among subpopulations, particularly young ones. For most genomic summary statistics, isolation from neighboring subpopulations was the most important factor driving variation, suggesting that, especially in remote parts of the metapopulation, there is moderate to low gene flow amongst subpopulations. In more isolated subpopulations, we found lower synonymous genomic diversity, π_S_,—a proxy for *N_e_*—(Figure 5), and in subpopulations with lower π_S_, we found higher rates of nonadaptive nonsynonymous substitutions, i.e., deleterious substitutions (Figure S3). Our evolutionary model of the *D. magna* metapopulation contrasts markedly with a much larger and older population of the same species, which was considerably more diverse and showed more efficient purifying selection, likely due to weaker genetic drift. These results confirm the theory of lower adaptive evolution in some metapopulations (Whitlock, 2004). By directly comparing the metapopulation and the stable population on a genomic level and using simulations, we can better understand the evolutionary dynamics in our focal metapopulation and how it is distinct from the evolution of large stable populations.

### Accumulation of deleterious mutations

It has been shown that *N_e_* explains cross-species variation in the rate of molecular evolution (Eyre-Walker, 2006; Eyre-Walker & Keightley, 2009; Gossmann et al., 2012). The within-species analog of this prediction is that in populations of different sizes, as they are commonly found in metapopulations, differences in genomic diversity are expected to explain variation in the rate of molecular evolution among (sub)populations. Populations with lower *N_e_* are expected to accumulate deleterious mutations faster, thus displaying higher *ω_NA_*. Consequently, their rate of adaptive substitution, *α*, would be lower, as shown across species by Galtier (2016). In our focal metapopulation, we found higher *ω_NA_* in subpopulations with smaller π_S_ (a proxy for lower *N_e_*; Figure S3). Consequently, *ω_A_* in subpopulations with smaller π_S_ was decreased, indicating that selection is less efficient in these subpopulations and that genetic drift may reduce the efficiency of selection. Plotting π_N_ against π_S_ shows a strong correlation, but with a slope clearly smaller than one (Figure 3). Thus, in subpopulations with lower π_S_, nonsynonymous diversity is higher relative to synonymous diversity than in subpopulations with higher π_S_ (Figure 3). This relationship might be because 1) purifying selection removes nonsynonymous deleterious mutations more efficiently in larger subpopulations, 2) gene flow masks genetic load, so that subpopulations with a lower π_N_/π_S_ ratio are less isolated (Figure 5). In the second scenario, recessive deleterious mutations could accumulate after a colonization bottleneck and subsequent inbreeding. Gene flow increases heterozygosity by introducing variation from less-related individuals, which would result in hybrid offspring with increased fitness, i.e., hybrid vigor (Ebert et al., 2002; Lohr & Haag, 2015).

Leroy et al. (2021) showed the effect of *N_e_* on (non-)synonymous diversity and (non-)adaptive evolution in insular versus continentally-distributed populations of different bird species. Their comparison also included an insular and a continental sample from the same species (*Fringilla coelebs*). *N_e_* was positively associated with *ω_A_*. We undertook a similar approach within a metapopulation of a single species with subpopulations of different ages and isolation levels and found the same relationship between π_S_ and *ω_A_*. This confirms that π_S_ explains variation in evolution within a species just as it does across species, as previous studies have noted (Eyre-Walker, 2006; Eyre-Walker & Keightley, 2009; Gossmann et al., 2012). By simulating different-sized populations in-silico to isolate the effects of *N_e_* on *ω_A_*, we further supported this finding, thereby obtaining the same associations between π_S_, π_N_/π_S_, and *ω* as we showed in the natural populations. Our results are also consistent with results by Fields et al. (2018), who showed that in genotypes collected from different *D. magna* metapopulations, protein-coding genes in the mitochondrial genome show enrichment in deleterious mutations compared to genotypes collected from larger and more stable populations in other parts of the species’ range.

### Age and isolation as predictors for genomic diversity

Once new subpopulations are founded by one or several individuals, they undergo rapid population expansion (clonal expansion in the case of cyclic parthenogens like *Daphnia*) and inbreeding as a consequence of sexual reproduction (Haag et al., 2005). This leads to low genomic diversity. Over time, immigration into subpopulations will increase genomic diversity. If subpopulations are inbred, the rate of effective gene flow may be elevated by hybrid vigor, as has been shown experimentally for the *D. magna* metapopulation (Ebert et al., 2002). In our focal metapopulation, we confirmed that genomic diversity increases with subpopulation age. We tested if this gain in genomic diversity is faster with a higher immigration rate, by quantifying the correlation between mean distance to the two closest neighboring subpopulations (NN2). As expected, isolated subpopulations have indeed lower genomic diversity than less isolated ones. Without further immigration, genomic diversity may even decrease rather than increase, as diversity may be lost by drift. There were, however, not sufficient isolated subpopulations that were old enough to test this assumption.

Earlier studies speculated that larger, more stable ponds with older subpopulations could be a reservoir for genomic diversity (Haag et al., 2005; Pajunen, 1986; Vanoverbeke et al., 2007); however, our analysis of the relationship between genomic diversity and the ecological characteristics of ponds does not support this suggestion. Larger ponds do not differ from other ponds in their genomic diversity. We also tested a finding from previous research in this metapopulation that infection with a virulent microsporidian parasite, which occurs in more than half of all subpopulations, is associated with genomic diversity (Cabalzar et al., 2019). In or study we do not observe this correlation. However, this earlier study focused exclusively on old subpopulations and aimed to understand the evolutionary differentiation of old subpopulations evolving with and without the parasite. Here, the inclusion of younger subpopulations prevented us to observe this correlation.

### Genomic differentiation is driven by geographical isolation

Genomic differentiation between subpopulations is generally high, mainly because of the high turnover dynamics in this metapopulation. These dynamics lead to frequent (re)colonization of vacant habitat patches, whereby the expectation of *F*_ST_ between newly founded populations is 0.5 (Wade and McCauley 1988). We estimated population differentiation between recently founded _subpopulations at ∼0.49, close to the theoretical expectation. However, as populations get older_ and receive immigrants, pairwise *F*_ST_ values were expected to decrease, which is what we observed.

For passively dispersed aquatic invertebrates, founder effects have been suggested as a main driver of differentiation; these include a combination of a few population founders, high population growth rates, and large population census sizes (Montero-Pau et al., 2018). Founder effects occur because many aquatic invertebrates are cyclic parthenogens, so a single individual can found and, after clonal expansion, populate the entire habitat patch. Montero-Pau et al. (2018) have shown that, in passively dispersed aquatic invertebrates, founder effects outweigh selective processes and migration. These factors were all considered equally important in the so-called monopolization hypothesis for explaining the genetic structure of aquatic invertebrates (De Meester et al., 2002). However, under the monopolization hypothesis, selective processes (e.g., adaptation) might be more efficient in populations with large *N_e_* that exhibit weaker genetic drift. Specifically, selection acts against the immigration of deleterious alleles and residential allele frequencies are favored (Lohr & Haag, 2015). Likewise, local adaptation, which is often observed in strongly structured aquatic populations (Decaestecker et al., 2007; Franch-Gras et al., 2017), may reduce the effective immigration rate of non-adapted genotypes, but much more so in populations with large *N_e_*. In the small subpopulations of our metapopulation, local adaptation has so far not been observed (Cabalzar et al., 2019; Roulin et al., 2015).

We also show that subpopulations separated by greater distance are more dissimilar than geographically closer ones, i.e., a pattern of IBD. On a local scale, previous metapopulation studies of freshwater zooplankton presented evidence against IBD (Martin et al., 2021; Montero-Pau et al., 2017). We found IBD both within and between islands, suggesting that gene flow is more likely between geographically closer subpopulations from the same and from different islands. This pattern is consistent with previous findings that subpopulations on the same island are genetically less differentiated than subpopulations on different islands (Roulin et al., 2016) and that dispersal distance exponentially decays, in which case long distance colonization events are rare (Dubart et al., 2020; Pajunen, 1986; Pajunen & Pajunen, 2003). As described for several aquatic organisms, sequential colonization can shape IBD (Montero-Pau et al., 2018). However, the long-term data on local extinction and re-colonization in this metapopulation suggest that the gene flow hypothesis might explain more of the observed pattern of IBD (Dubart et al., 2020; Pajunen, 1986; Pajunen & Pajunen, 2003). The isolation measure, NN2, was the variable that best explained a subpopulation’s *F*_ST_ estimated with the *F*-model approach and could be shown to correlate with pairwise *F*_ST_ (Figure 7). Finding an association between NN2 and *F*_ST_ in these two complementary approaches underlines the importance of gene flow in predicting genomic differentiation in this metapopulation (Gaggiotti & Foll, 2010). Our population structure analysis agrees with a pattern of IBD.

## Conclusion

We studied the population genomics of a well-documented, highly dynamic metapopulation to understand if metapopulations evolve differently from panmictic populations. The obvious differences between these two types of populations are that metapopulations feature extinction– (re)colonization dynamics and gene flow. The genomic consequences of metapopulation ecology include recurrent bottlenecks during population founding, which can lead to high genomic differentiation between subpopulations and low genomic diversity within subpopulations. Even though *D. magna* census population sizes may be large, bottlenecks cause low genomic diversity and strong genetic drift, which in turn reduces the efficacy of selection. This is seen as lowered rates of adaptive nonsynonymous substitutions and elevated numbers of nonsynonymous relative to synonymous polymorphisms in the metapopulation compared to the single, large, stable population. Our observed and simulated results support the expected differences between these two population types and suggest that nondeterministic forces dominate the evolutionary process in dynamic metapopulations. Our study does not only provide genomic insights into a well-documented metapopulation, but we also link genomics to subpopulation ecology and thus, unravel the evolutionary mechanisms of a metapopulation in a fragmented habitat. This provides valuable insights for conservation biology and helps to understand how metapopulations evolve differently from Wright–Fisher populations. Our findings are largely consistent with the propagule model of metapopulation evolution (Slatkin, 1977).

## Acknowledgments

We thank V. Ilmari Pajunen, Irmeli Pajunen, Christoph R. Haag, Jürgen Hottinger, Frida Ben-Ami, Andrea Cabalzar, Mikko Lehto, Jennifer Lohr, David Preiswerk, David Duneau, Katharina Ida Ebert, Gleb Georg Ebert, A. Marcelino, Y. Haag, E. Haag, C. Liautard-Haag, C. Reisser, C. Molinier, E. Hürlimann, and C. Mills for help in the field. We thank members of the Ebert group for feedback on the study and the manuscript. This work was supported by the Swiss National Science Foundation (SNSF) (grant numbers 310030B_166677, 310030_188887 to DE).

## Data accessibility

Analysis scripts are available at https://github.com/pascalangst/Angst_etal_2022_XXX, and raw data are deposited at the NCBI SRA database (BioProject IDs XXXX, YYYY).

## Author contributions

PA, DE, and PDF designed the study. PA and PDF analyzed the data. DE supervised the long-term metapopulation data collection and curates the data. CA collected and prepared the Aegelsee sample. DE and PDF collected, and PA and PDF prepared the metapopulation samples. PA wrote the manuscript. All authors reviewed the manuscript.

## Notes

### Competing Interest Statement

The authors have declared no competing interest.

