## Supplemental Material for "Genetic drift shapes the evolution of a highly dynamic metapopulation"

**Supplemental Results**

### D. sinensis genome annotation

Annotation of the *D. sinensis* genome resulted in nearly 21,000 gene models. The observed number of gene models in *D. sinensis* is lower than the number of gene models in the focal species, *D. magna*, though the number is not so different from the number of gene models found in other *Daphnia* species such as *D. pulex* (around 19 thousand; Ye et al., 2017). A possible explanation for the lower number of annotated genes is the more limited RNAseq resources presently available for *D. sinensis* as compared to *D. magna*. However, the number of genes present in this *de novo* annotation, and in particular the number of identifiable orthologs between the species, is more than sufficient for the target analysis of a genome-wide summary of selection on protein-coding regions.

**Figure S1: Dimensionality-reduction using PCA and t-SNE.** This is the same figure as Figure 2 but with numbers added to ponds. PCA (A) and t-SNE (B) based on whole-genome allele frequency data reveal spatial population structure. Samples from the same island form clusters in most cases (see Figure 1). A combination of colors and symbols indicates each island of origin. The ponds on the same island are numbered consecutively.

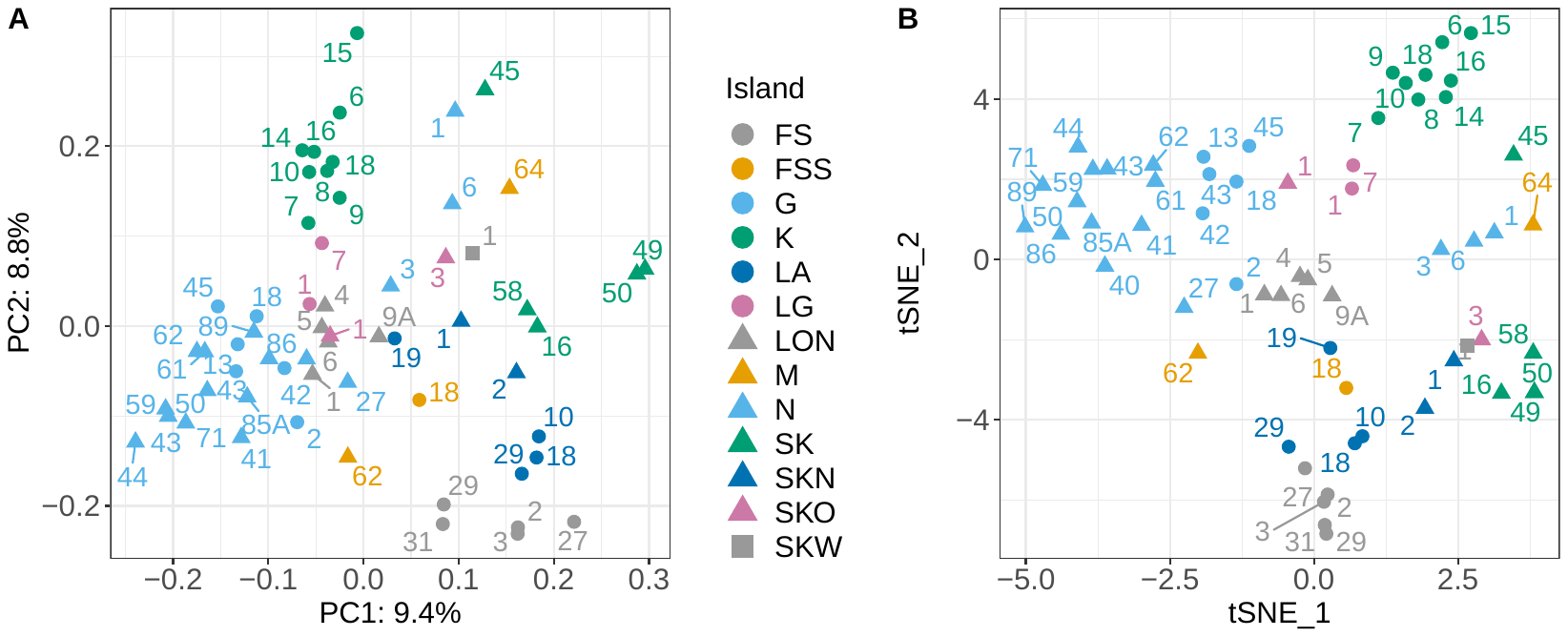

**Figure S2: Association between (non-)synonymous genomic diversity and *N_e_*.** This is the same figure as Figure 3A but with numbers added to ponds. Observed π_N_ and π_S_ in the metapopulation. More diverse subpopulations have relatively fewer nonsynonymous polymorphisms. The color, symbol, and number scheme are the same as in Figure S1. The size of this symbol is indicative for the subpopulation’s NN2. The black line indicates the one-to-one ratio.

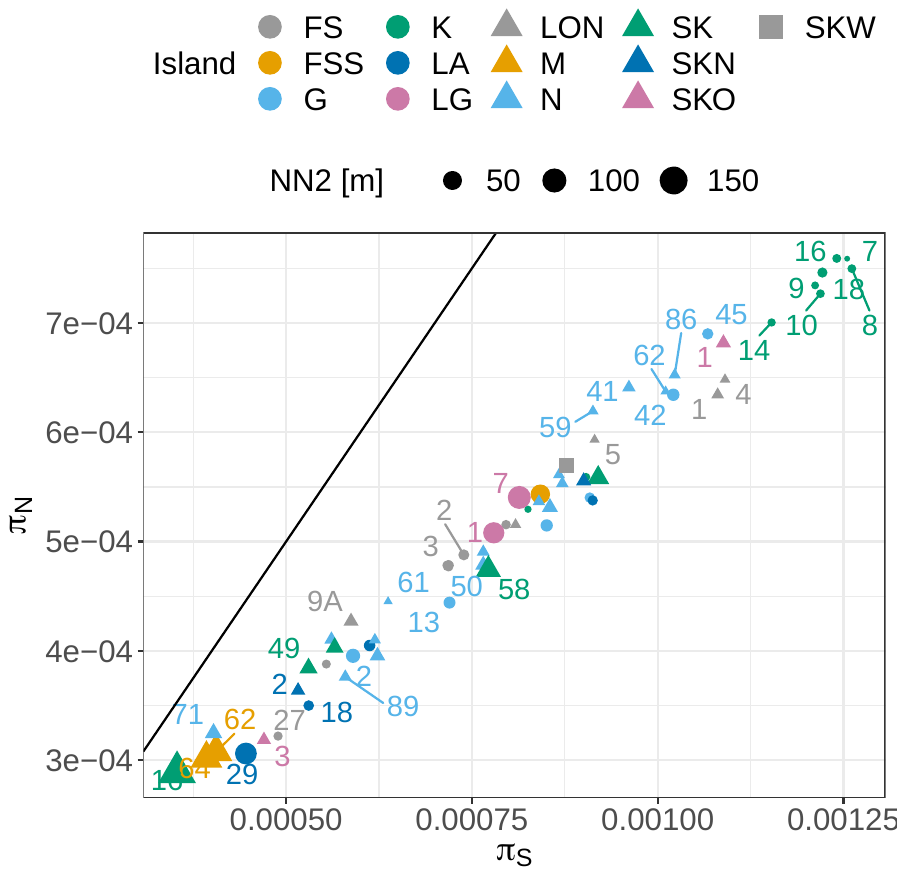

**Figure S3: Association between nonadaptive nonsynonymous substitutions and π_S_.** Depicted is observed *ω_NA_* of each subpopulation of the metapopulation. *ω_NA_* is negatively associated with π_S_. π_S_ can be used as an approximation for *N_e_*. The red dot is the estimate for the single, large, stable population from Switzerland.

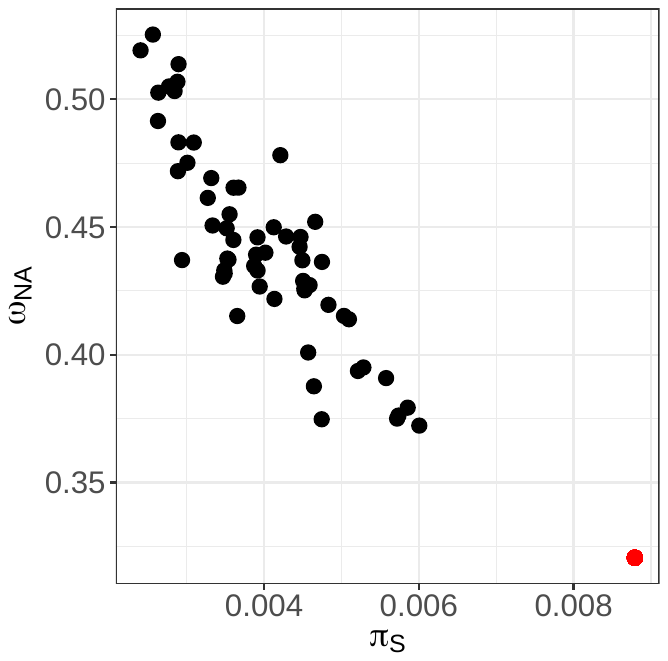

**Figure S4: Simulated rate of nonadaptive nonsynonymous substitutions in differently sized populations.** *ω_NA_* is negatively associated with synonymous heterozygosity, *H_S_*, and simulated population size. *H_S_* can also be used as an approximation for *N_e_*. Horizontal and vertical error bars indicate the standard error around the mean of 1,000 runs (simulations were made using SLiM).

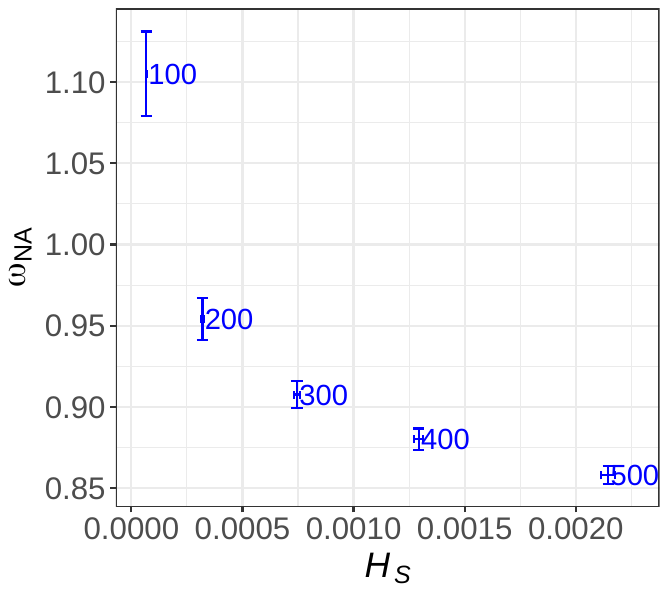

**Figure S5: Genomic diversity explained by subpopulation age and isolation.** This is the same figure as Figure 6 but with numbers added to ponds. Genomic diversity, π, is positively correlated with subpopulation age and negatively correlated with the isolation measure, NN2. The subpopulation age was added to one before log_10_-transformation. The color, symbol, and number scheme are the same as in Figure S1. The size of this symbol is indicative for the subpopulation’s NN2. The black line is a regression line based on a linear model with the confidence interval depicted as shading around the line.

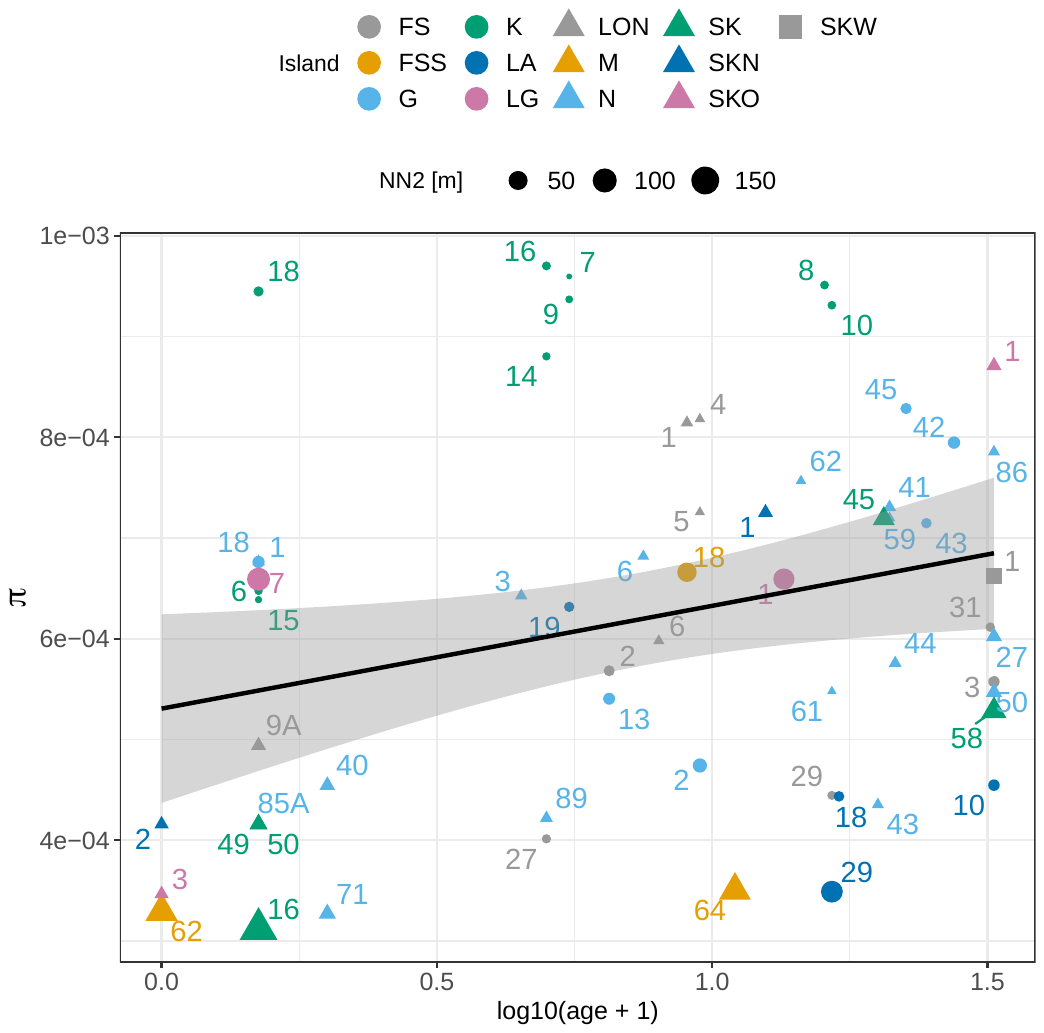

**Figure S6: Isolation by distance (IBD).** This is the same figure as Figure 7 but with numbers added to ponds. *F*-model-based *F*_ST_ positively correlates with the isolation measure NN2 [m]. Statistics derive from a Spearman correlation test. The color, symbol, and number scheme are the same as in Figure S1.

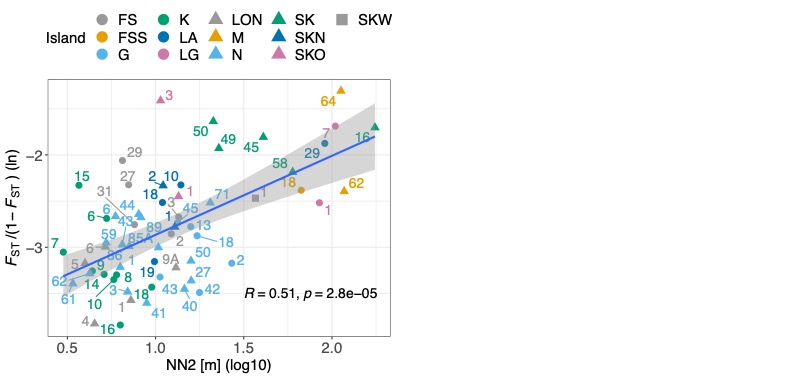

**Figure S7: Isolation by distance (IBD).** Pairwise population differentiation is positively correlated with geographic isolation measured as NN2. Each point represents the comparison between two subpopulations. Statistics derive from dbMEM analysis by RDA. The blue line is the regression line of a linear model with the 95 % confidence interval depicted as shading around the line.

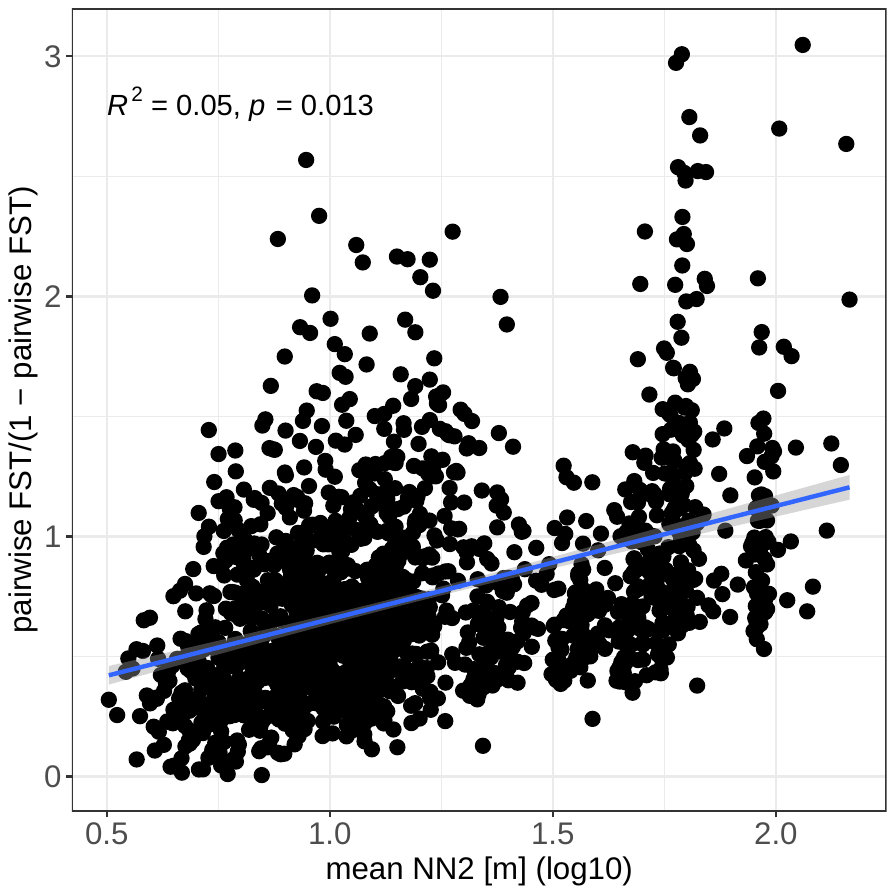

**Table S1: Detailed sample information and mapping statistics.** Each subpopulation sample (Pond ID) is accompanied by the number of pooled animals prior to DNA extraction (#animals), the percentage of sequencing reads that mapped to the *D. magna* reference genome (mapping_percent), the average whole-genome coverage (average_coverage), and the infection status with *H. tvaerminnensis* (infection_status).

| **Pond ID** | **#animals** | **mapping_percent** | **average_coverage** | **infection_status** |
| --- | --- | --- | --- | --- |
| FS-2 | 9 | 0.9886 | 84.383 | 0 |
| FS-27 | 27 | 0.9399 | 22.6618 | 1 |
| FS-29 | 17 | 0.9925 | 50.4757 | 0 |
| FS-3 | 11 | 0.9649 | 67.5036 | 1 |
| FS-31 | 31 | 0.918 | 53.9897 | 1 |
| FSS-18 | 52 | 0.9886 | 76.6795 | 1 |
| G-13 | 38 | 0.8331 | 19.9154 | 1 |
| G-18 | 51 | 0.9872 | 66.0887 | 0 |
| G-2 | 19 | 0.9721 | 62.5149 | 0 |
| G-33 | 39 | 0.9799 | 0.321314 | NA |
| G-42 | 51 | 0.9838 | 56.1439 | 1 |
| G-43 | 50 | 0.9507 | 23.5622 | 1 |
| G-45 | 44 | 0.9876 | 55.301 | 0 |
| K-10 | 28 | 0.8706 | 45.5364 | 1 |
| K-14 | 66 | 0.9432 | 48.357 | 1 |
| K-15 | 67 | 0.8737 | 30.4633 | 1 |
| K-16 | 54 | 0.8821 | 59.2523 | 1 |
| K-18 | 61 | 0.8831 | 64.912 | 1 |
| K-6 | 53 | 0.9642 | 21.4087 | 1 |
| K-7 | 71 | 0.9766 | 104.367 | 1 |
| K-8 | 53 | 0.8848 | 62.1974 | 1 |
| K-9 | 58 | 0.9571 | 50.9035 | 1 |
| LA-10 | 28 | 0.9823 | 21.1503 | 1 |
| LA-18 | 30 | 0.9877 | 20.8245 | 1 |
| LA-19 | 71 | 0.9845 | 21.7759 | 1 |
| LA-29 | 59 | 0.9804 | 24.0561 | 0 |
| LG-1 | 40 | 0.9873 | 64.8407 | 1 |
| LG-7 | 49 | 0.9889 | 43.4276 | 0 |
| LON-1 | 58 | 0.8055 | 37.0342 | 1 |
| LON-4 | 55 | 0.7394 | 31.6327 | 1 |
| LON-5 | 53 | 0.9572 | 48.7995 | 1 |
| LON-6 | 51 | 0.9094 | 20.9864 | 1 |
| LON-9A | 14 | 0.955 | 25.4082 | 1 |
| M-62 | 50 | 0.948 | 24.7763 | 1 |
| M-64 | 42 | 0.9871 | 47.523 | 0 |
| N-1 | 11 | 0.9729 | 46.767 | 1 |
| N-27 | 49 | 0.9625 | 20.2174 | 1 |
| N-3 | 56 | 0.931 | 18.6522 | 1 |
| N-40 | 46 | 0.8756 | 18.6395 | 1 |
| N-41 | 49 | 0.9925 | 61.2961 | 0 |
| N-43 | 53 | 0.9034 | 16.1057 | 1 |
| N-44 | 49 | 0.938 | 23.3142 | 1 |
| N-50 | 54 | 0.9432 | 21.2262 | 1 |
| N-59 | 36 | 0.8319 | 54.2554 | 1 |
| N-6 | 53 | 0.9571 | 36.5148 | 1 |
| N-61 | 37 | 0.7418 | 17.5543 | 1 |
| N-62 | 57 | 0.9247 | 36.0808 | 1 |
| N-71 | 54 | 0.7249 | 15.3076 | 1 |
| N-85A | 54 | 0.8932 | 18.5148 | 1 |
| N-86 | 49 | 0.979 | 51.3107 | 1 |
| N-89 | 50 | 0.8405 | 30.9543 | 1 |
| SK-1 | 61 | 0.9458 | 0.230235 | NA |
| SK-16 | 60 | 0.9866 | 20.1261 | 0 |
| SK-45 | 37 | 0.8716 | 43.4562 | 1 |
| SK-49 | 50 | 0.9935 | 27.6059 | 0 |
| SK-50 | 48 | 0.9888 | 47.9354 | 0 |
| SK-58 | 48 | 0.9683 | 19.4414 | 1 |
| SKN-1 | 49 | 0.9006 | 52.9265 | 1 |
| SKN-2 | 61 | 0.9744 | 21.7282 | 1 |
| SKO-1 | 45 | 0.9706 | 57.1013 | 1 |
| SKO-3 | 51 | 0.9883 | 37.4972 | 0 |
| SKW-1 | 50 | 0.9862 | 23.9365 | 1 |

**Table S2: Type two analysis of variance between genomic diversity, π, and explanatory variables, including the island of origin.** After including the island of origin, the association to the measure of isolation became weaker and insignificant, since isolated subpopulations are mostly from the same, isolated islands (*F*(12,47) = 13.57, *p* < 0.001, *R*^2^ = 0.776). One *R*^2^ value is given for the complete model. Significant associations are in bold.

| Explanatory variable | df | Statistics | *p* | *R*^2^ |
| --- | --- | --- | --- | --- |
| Population age | 42 | t = 2.712 | **0.0004** | 0.673 |
| PC1 (marineness) | 42 | t = -0.822 | 0.642 |  |
| PC2 (pond geometry) | 42 | t = 0.259 | 0.874 |  |
| NN2 (isolation) | 42 | t = -1.538 | 0.060 |  |
| Island | 12/42 | F = 5.0021 | **4.4 × 10^-5^** |  |
| Infection status | 1/42 | F = 0.1464 | 0.704 |  |
